# Human–Dog Interaction Method and Dog Familiarity Differentially Modulate Prefrontal Connectivity and Autonomic Recovery Following Acute Stress: An fNIRS Study

**DOI:** 10.64898/2026.03.25.714338

**Authors:** Brianna Kish, Reina Nishiura, Niwako Ogata, Yunjie Tong

## Abstract

Human–dog interaction is widely used to alleviate stress, yet the accompanying cortical and autonomic dynamics during acute stress and recovery remain incompletely characterized. In this study, 70 adult dog owners completed a standardized stress protocol while prefrontal cortex activity was continuously monitored with functional near-infrared spectroscopy (fNIRS), alongside subjective stress and salivary cortisol measures. Participants then underwent a recovery phase involving interaction with a companion dog, manipulating contact type (direct in person vs. indirect video conferencing), and familiarity (own vs. unfamiliar dog). Stress responses were quantified through heart rate (HR), heart rate variability (HRV), low- and high-frequency spectral power (LF, HF, and LF/HF), and prefrontal functional connectivity (FC) based on maximum cross-correlation coefficients between fNIRS channels. As expected, HR, HRV-derived indices, and FC increased from baseline to the stress phase, confirming robust engagement of autonomic and prefrontal networks. During the recovery phase, all dog interaction conditions demonstrated reductions in HR, LF/HF ratio, and FC toward or below baseline, consistent with physiological and neural stress recovery; direct interaction was associated with particularly pronounced parasympathetic enhancement and a drop in FC that fell significantly below baseline in some cases. Across groups, HRV, LF/HF, and FC were the most consistent predictors of subjective stress ratings, whereas associations with cortisol were limited. These findings suggest that human–dog interaction promotes coordinated autonomic and prefrontal recovery from acute stress, and that fNIRS-derived metrics might provide a marker of stress modulation that can distinguish high-cognitive load and low cognitive demand states beyond traditional stress indices.

## Introduction

The positive effects of human–animal interactions on stress regulation, mood, and overall well-being have been widely documented. Such interactions have been associated with improvements in both physical and mental health, including cognitive function and emotional stability ^1–3^. In recent years, companion animals and human-animal interactions have gained increasing prominence in healthcare and quality-of-life programs, particularly for individuals with mood-related disorders such as post-traumatic stress disorder and depression ^4,5^. Among companion animals, dogs are most commonly employed, with human–dog interactions demonstrating a range of physiological benefits, including reduced heart rate and blood pressure, as well as psychological benefits such as decreased loneliness and improved mood ^1,2,6,7^.

Although the long-term health benefits of animal companionship are well established, such as the reduction in risk of developing hypertension and improved blood pressure control^8^, the short-term effects of human–dog interactions, particularly as mediators in stressful environments, remain less clearly defined. Studies have consistently shown that interaction with animals can reduce perceived stress; however, the underlying mechanisms of this effect are not yet fully understood ^9–20^. For example, Allen et al. reported significantly lower heart rate and blood pressure in participants undergoing stress testing when a dog was present ^21^. Similarly, Martos-Montes et al. observed rGeduced heart rate, but no change in blood pressure, among university students completing the Trier Social Stress Test in the presence of a dog ^22^. In contrast, Machová et al. found that a brief, 10-minute dog interaction improved mood without corresponding changes in heart rate or blood pressure ^23^. These inconsistent physiological findings suggest that traditional measures alone may be insufficient to fully characterize the stress response and recovery process.

Neuroimaging techniques offer new opportunities to examine the psychological and neural mechanisms underlying human–animal interactions. Studies using functional near-infrared spectroscopy (fNIRS) have reported increased prefrontal cortex activation during interaction with dogs compared to rest ^24,25^, while electroencephalography research has demonstrated increased alpha and beta power, indicating enhanced concentration without stress ^26^. Additionally, Lee et al. found that even indirect exposure to dogs, such as watching videos of dogs playing, produced similar alpha and beta power increases ^27^. These findings suggest that dog-related stimuli may assist in improved stress response barring direct physical contact.

Despite these advances, important gaps remain in the literature. Few neuroimaging studies have examined the role of human–animal interaction in stress recovery, while considering individual differences such as the degree of animal familiarity, or directly comparing different modes of interaction. This limitation is particularly relevant in the modern day, where pet owners increasingly interact with their animals digitally through platforms such as pet cameras and video calls ^28^.

Understanding whether indirect interactions provide comparable stress-relieving effects to in-person contact is therefore of growing importance.

To address these gaps, the present study employs fNIRS to investigate the physiological, psychological, and neural correlates of human–dog interaction during stress. Specifically, we aim to answer the following questions:

1. How does dog interaction mediate the stress response at both physiological and psychological levels?
2. Does the mode of interaction (direct versus indirect, familiar versus unfamiliar dog) modulate the stress response?
3. Do fNIRS-derived measures of neural activity correlate with perceived stress?

## Methods

### Participants

70 dog-owner pairs were successfully recruited for this study (male: 5, female: 65, non-binary/third gender: 1; aged 18-35 yrs: 43, aged 36-55 yrs: 28). Participants were recruited under the following criteria. Owners must: be aged 18-55 years, have been their dog’s companion for 6+ months, possess no circulatory or panic disorders, and not take any specific medications. Their dogs must: be aged 1-12 years, weigh over 15 lbs., have no major health problems affecting activity, and be friendly and calm (i.e., do not exhibit aggressive behavior or severe separation anxiety). Written informed consent was obtained from all participants.

### Experimental Protocol

Demographic information was collected before the first visit via an online survey (see Figure 1a for full list). During each visit, participants were fitted with a functional near-infrared spectroscopy (fNIRS) cap consisting of 20 channels covering the prefrontal cortex (sampling rate: 7.185 Hz; probe layout shown in Figure 1b; NIRScoutXP, NIRx Medizintechnik GmbH, Berlin, Germany). fNIRS was chosen to monitor neuronal engagement for this experiment as it measures changes in oxy-/deoxy-genated hemoglobin. Through neurovascular coupling, this allows us to see both neuronal demand and changes in blood flow due to basic physiology, such as heartbeat. A baseline salivary cortisol sample (higher = greater physiological stress) and subjective stress rating, assessed using a visual analogue scale (VAS, range: 0-100, higher = greater subjective stress), were then collected.

**Figure 1:**
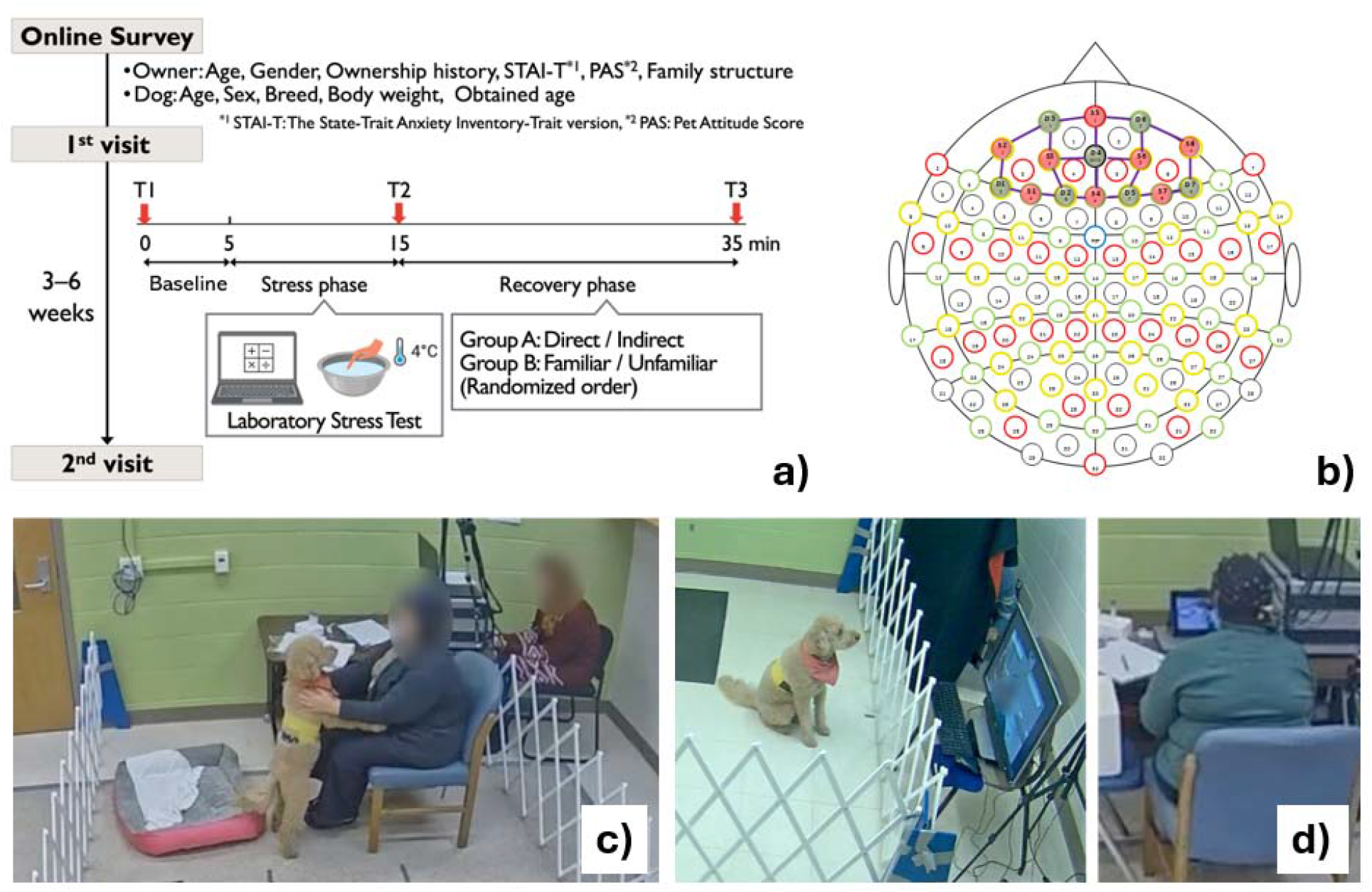
Experimental Setup. A) Experimental protocol. Participants completed an online demographic survey prior to the 1^st^ visit. During both visits, fNIRS was collected during a 3 -min baseline scan, 10-min stress test, and 20-min recovery phase. Salivary cortisol samples and subjective stress scores were collected at timepoints T1, T2, and T3. Group A participants interacted with their dog directly or indirectly. Group B participants interacted with their own dog or an unfamiliar dog indirectly. Participants returned for their 2^nd^ visit to repeat the process with the alternative recovery phase condition. B) fNIRS probe configuration, 7 detectors (green) and 8 sources (red) were placed across the prefrontal cortex (20 channels). C) Direct dog condition example. D) Indirect dog condition example.

Participants then completed a 3-minute resting-state fNIRS recording, during which they were instructed to remain still and focus on a visual fixation cross. This was followed by the completion of the Maastricht Acute Stress Test (MAST). This stress task employs an alternating block design consisting of a cognitive stressor (mental arithmetic) and a physical stressor (cold pressor test), in which participants submerged their left hand in ice water (4 °C), for a total duration of ∼10 minutes ^29^. Following the stress task, post-stress salivary cortisol samples and VAS rating were obtained.

Participants subsequently began a 20-minute recovery phase and were assigned to one of two groups based on the type of dog interaction. Group A (n = 40) engaged in either direct or indirect interaction with their own dog (Figure 1c), whereas Group B (n = 30) engaged in indirect interaction with either their own familiar dog or an unfamiliar dog (Figure 1d). The order of interaction conditions was randomized across visits for each participant.

In the direct interaction condition, participants’ dogs were allowed to freely roam in a pen alongside their owner. Participants were instructed to interact with their dogs in any manner they felt most stress-relieving (e.g., petting, watching, speaking), while minimizing head movement to reduce motion artifacts in the fNIRS signal (Figure 1c). In the indirect interaction condition, dogs were placed in a separate room and interacted with participants via audio-visual communication (Zoom, Zoom Video Communications, Inc., San Jose, California, USA). Participants could also remotely give treats to their dogs through verbal request (Figure 1d). Following the recovery phase, a final salivary cortisol sample and VAS rating were collected.

### Data Analysis

All fNIRS data were preprocessed in MATLAB (MATLAB 2021b; The MathWorks Inc., Natick, MA, 2000) with the following pipeline: conversion from light absorption to change in oxy- and deoxy-hemoglobin concentration (Δ[HbO] and Δ[Hb] respectively), motion correction (kurtosis-based wavelet algorithm)^30^, detrending, and bandpass filtering (0.01-2 Hz). Within individual subjects, channels with excessive motion or poor signal quality (e.g., no heartbeat detected) were excluded from analysis. 2 subjects’ scans from Group A and 2 scans from Group B were excluded due to poor data quality. Δ[HbO] signals were used for all further analyses due to increased signal strength.

The data was then split into the 3-min baseline, 10-min stress test, and 20-min relaxation phase. Data from the relaxation phase was further split into the first and last 10 minutes, as human behavior is likely to fluctuate over long time periods. To assess the physiological stress response, heart rate (HR), heart rate variability (HRV), total low-frequency power (LF; 0.04-0.15 Hz), total high-frequency power (HF; 0.15-0.4 Hz), and the ratio of low-frequency power to high-frequency power (LF/HF) were calculated for each phase. HR is reported as a change in HR from baseline measurements to account for individual variability in base HR. HRV was calculated as the ratio of the standard deviation of inter-beat intervals (SDNN) to the root mean squared of the derivative of inter-beat intervals (RMSSD), as this method reflects the balance between sympathetic and parasympathetic nervous system activity, with a higher ratio indicating sympathetic dominance ^31^. Spectral power was determined through the MATLAB functions pwelch and bandpower. Lastly, the psychological stress response was measured through functional connectivity (FC). In brief, the fNIRS time series were filtered to the neuronal low-frequency range (0.01-0.1 Hz), and cross-correlation was run between all channel pairs. The average maximum cross-correlation coefficient (MCCC) across all channel pairs was reported as FC. MCCCs under 0.3 were excluded before averaging to remove spurious correlations ^32^.

### Statistical Analysis

4-way analyses of variances (ANOVA) were computed for all fNIRS-derived metrics to compare across experimental phases: Baseline, Stress, and Relaxation (Group A: Indirect or Direct, Group B: Familiar or Unfamiliar), accounting for age, gender, and baseline VAS. Baseline VAS was included to account for the initial stress levels of participants prior to the start of the experimental paradigm. Multiple comparisons between groups using Tukey’s honest significant test were then performed to determine significance. Additionally, linear modeling was run between the fNIRS metrics and cortisol and VAS data, adjusting for age and gender. One subject from Group A was excluded from linear modeling due to invalid cortisol values.

## Results

Figure 1a and 1b present the results of the HR analysis. HR was shown to increase from baseline to the stress phase and then drop significantly during the relaxation phase, aligning with prior research using the MAST ^33^. All interaction conditions showed significant decreases with no difference between direct and indirect for Group A or familiar and unfamiliar for Group B. In HRV, significant increases from baseline to stress are seen followed by a decrease during the relaxation phase. In Group A, this drop is most apparent in the 2^nd^ half of the relaxation phase. No significant differences were seen between interaction conditions.

LF power demonstrated no significant changes between experiment phase or interaction conditions for both group A and B. However, all phases show a non-significant increase from baseline (Figure 2a, b). HF power showed a similar trend in increasing power from baseline for both groups, and in group A, a significant increase in HF power is seen for the 1^st^ and 2^nd^ half direct interaction condition from baseline recordings (Figure 2c, d). Finally, for the LF/HF ratio, significant increases were seen from baseline to stress for both groups. In group A, a significant drop from the stress phase is only seen in the 2^nd^ half direct interaction condition (Figure 2e). In group B, significant decreases from stress are seen in all interaction conditions, with higher power in the 2^nd^ half; however, no differences between interaction conditions are present (Figure 2f).

**Figure 2:**
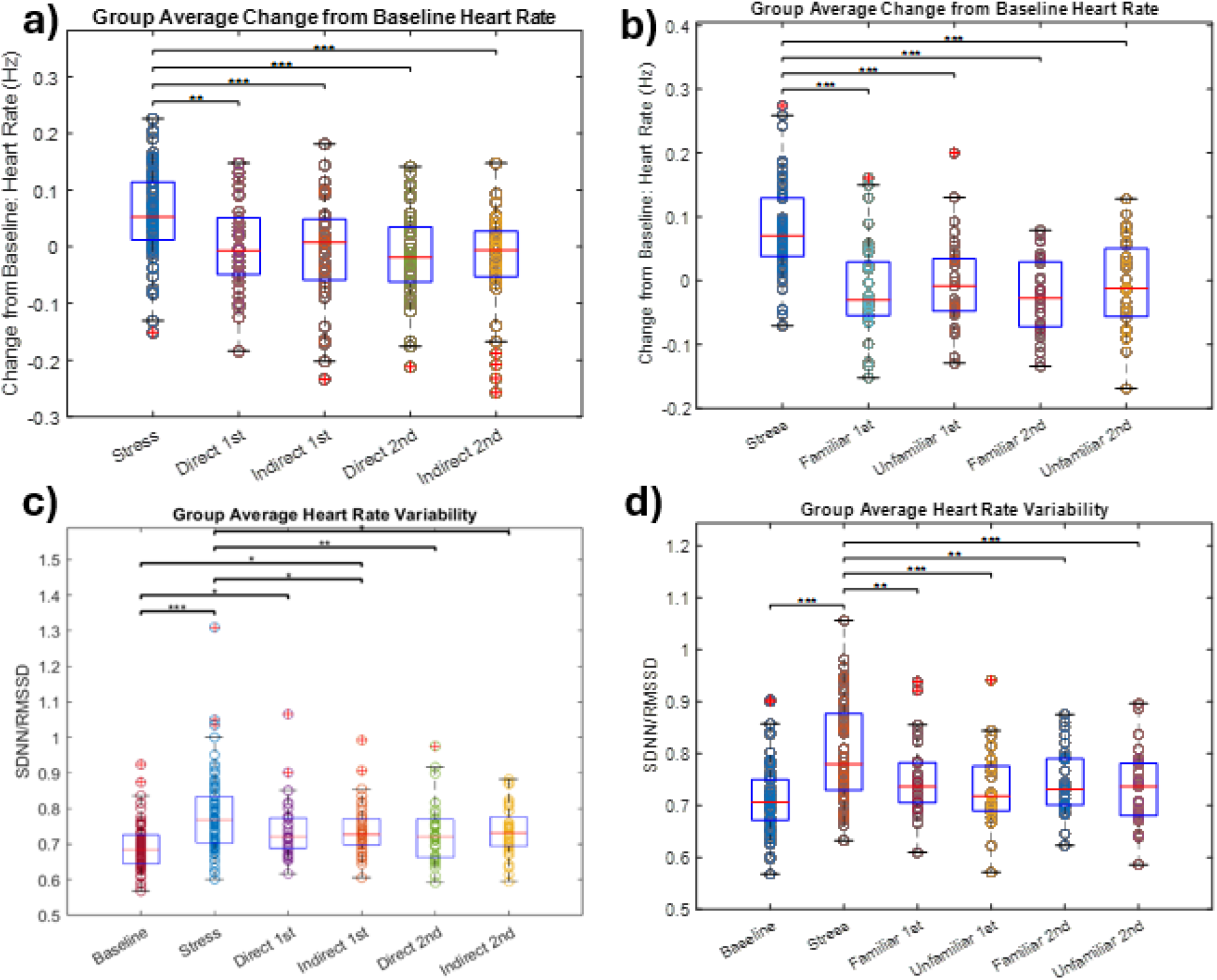
Heart Rate and Heart Rate Variability Changes during Stress Testing (Group A: left column, Group B: right column). A, B) Change from heart rate measured at baseline across stress and relaxation phases. C, D) Heart rate variability measured at baseline, stress, and relaxation phases. Relaxation phase split into 1^st^ and 2^nd^ 10-minute periods. Asterisks indicate significant differences between groups (*: p<0.05, **: p<0.01, ***: p<0.001)

In Figure 4, the results of functional connectivity analyses are presented. In group A, we see a significant increase in average MCCCs from baseline to stress, followed by a significant drop for all interaction conditions. Furthermore, MCCCs under the direct condition were significantly lower than baseline. This is not seen in the indirect condition, but no significant differences between direct and indirect were observed (Figure 4a). In Figure 4b, we see the same increase from baseline to stress, followed by decreases for all interaction conditions. No significant differences were found between baseline and interaction conditions or between the familiar and unfamiliar interaction conditions.

The results of linear regression between the 6 fNIRS-derived metrics and experimental data, cortisol and VAS, are reported in Table 1. Significant relationships (p < 0.05) are noted with an asterisk. In group A, we see that FC was a significant predictor of both cortisol and VAS, and HR, HRV, and LF/HF predictors of VAS. In group B, HRV, LF, HF, LF/HF, and FC were all significant predictors of VAS. No significant relationships were seen for cortisol. Across both groups, only HRV, LF/HF, and FC were significant predictors of VAS.

**Table 1:** Linear Modeling of fNIRS-Metrics with Cortisol and VAS. P-values are reported for linear regression of fNIRS-metrics: HR, HRV, LF, HF, LF/HF, and FC, to salivary cortisol and VAS data. Results adjusted for age and gender.

|  | **Group A** |  | **Group B** |  |
| --- | --- | --- | --- | --- |
|  | **Cortisol** | **VAS** | **Cortisol** | **VAS** |
| **HR** | 0.6939 | 1.0961e-04* | 0.2666 | 0.2523 |
| **HRV** | 0.3766 | 1.1044e-04* | 0.9155 | 0.0084* |
| **LF** | 0.9219 | 0.9421 | 0.0961 | 0.0323* |
| **HF** | 0.9241 | 0.9501 | 0.0634 | 0.0160* |
| **LF/HF** | 0.1290 | 0.0085* | 0.6002 | 0.0434* |
| **FC** | 0.0448* | 3.1643e-07* | 0.1923 | 0.0348* |

## Discussion

This study employed fNIRS to examine the psychological and physiological effects of stress on the human body and how it recovers during a human-dog interaction. Two modes of dog interaction were employed: direct versus indirect, and unfamiliar dog versus familiar dog. Our results indicate that most fNIRS measures increased under increased cognitive load and perceived stress triggered by a stress test before decreasing with the introduction of a dog companion. While no strong differences were found between unfamiliar and familiar dog effects, direct interaction over indirect appeared to influence stress recovery.

### Cardiac Measures

Both HR increased from baseline to stress before decreasing during the relaxation phase in groups A and B (Figure 2a, b). As part of the body’s “fight or flight” response, i.e., the sympathetic nervous system, the brain releases adrenaline in response to external stressors, increasing heart rate. As the body adapts to or removes the stressor, the “rest and digest” or parasympathetic nervous system activates, releasing acetylcholine, which decreases vagus nerve stimulation, and thus, heart rate. As seen in our results, the MAST was sufficient to drive a significant stress response, consistent with previous work, and dog interaction, enough to bring heart rate back to baseline ^34^. These fluctuations in HR in turn caused corresponding changes in HRV (Figure 2c, d). Calculating HRV as the ratio of SDNN to RMSSD highlights the effects of sympathetic to parasympathetic nervous system activity ^31^. Thus, our data aligns well with the known increase of sympathetic activity when exposed to stressful environments. It should be noted that all dog interaction conditions elicited a similar relaxation response, with no significant differences.

### Frequency Response

We further examined autonomic nervous system activity through the signals’ frequency content. Prior research has identified the low-frequency band (0.04-0.15 Hz) in blood-based signals as the component of HRV related to sympathetic nervous system activity, and complementarily, the high-frequency band (0.15-0.4 Hz) as the parasympathetic nervous system ^35^. Surprisingly, no significant differences across experimental phases were observed in LF power for both groups A and B; however, a trend of increase compared to baseline was present. In HF power, no change was seen in group B, but in group A, a significant increase from baseline to the direct interaction condition was found (Figure 3). This could indicate that direct dog interaction triggers significant parasympathetic activation, aligning with a stronger “rest and digest” function or calming effect. This is the first study of our knowledge to observe this comparison.

**Figure 3:**
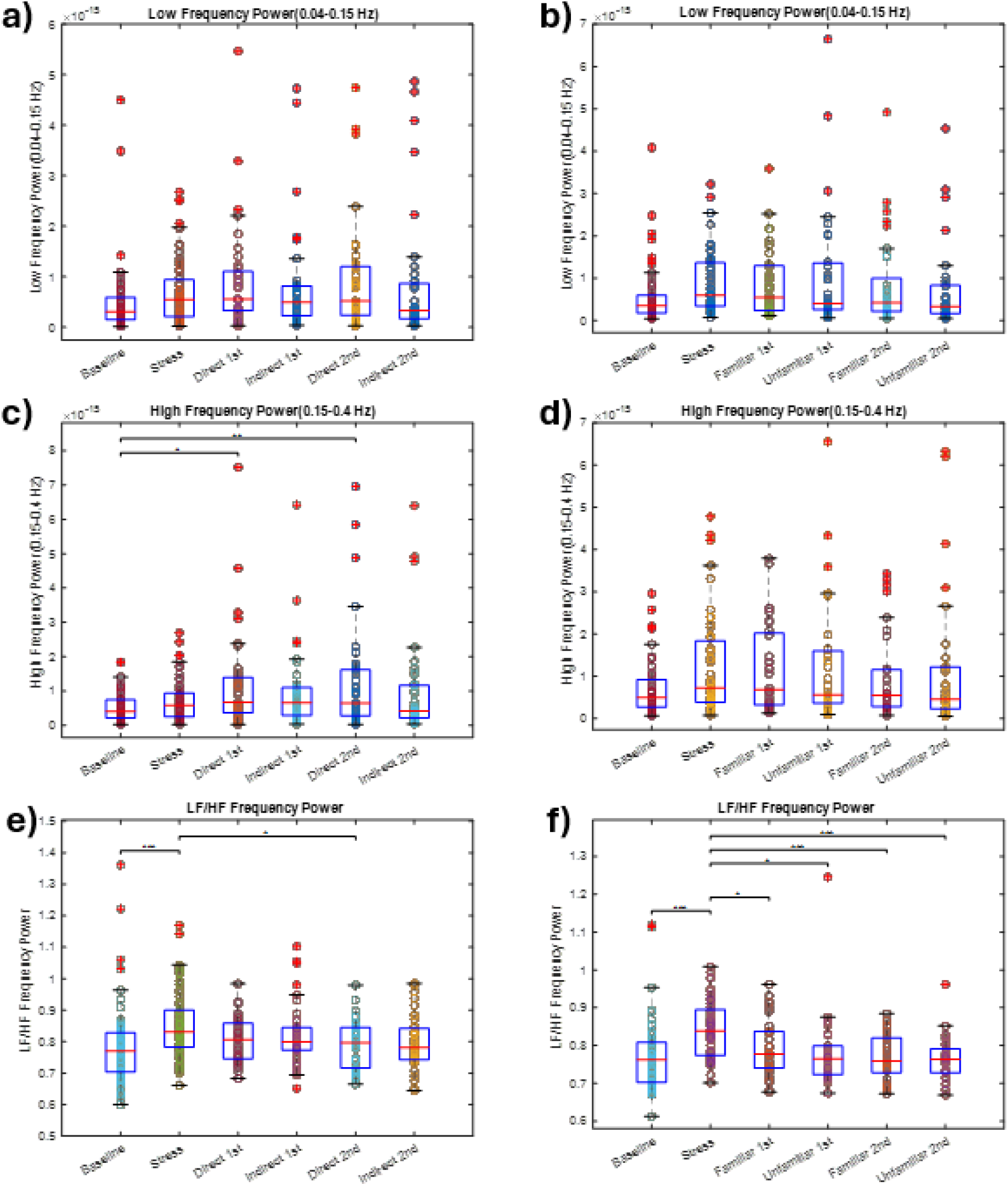
Total Frequency Bandpower during Stress Testing (Group A: left column, Group B: right column). A, B) Low-frequency (0.04-0.15 Hz) power measured at baseline across stress and relaxation phases. C, D) High-frequency (0.15-0.4 Hz) power measured at baseline, stress, and relaxation phases. E, F) Ratio of low-frequency power to high-frequency power measured at baseline, stress, and relaxation phases. Relaxation phase split into 1^st^ and 2^nd^ 10-minute periods. Asterisks indicate significant differences between groups (*: p<0.05, **: p<0.01, ***: p<0.001)

Similar to our method of calculating HRV, the LF/HF power ratio highlights sympathetic activity relative to parasympathetic activity. The results follow, with an increase from baseline to stress, then a decrease during the dog interaction phase. Interestingly, only the direct condition in group A showed a significant decrease from stress, while all conditions in group B were significantly lower (Figure 3e, f). Together, these results might indicate that sympathetic activity might not be the driving force in the changing physiology during our stress test, but rather the combined effects of sympathetic and parasympathetic activity instead.

### Neuronal Activity

Functional connectivity characterizes the coherence in activity of different brain regions. The prefrontal cortex (PFC) performs as the executive hub of information processing, decision making, and self-monitoring. Within the PFC, coherence and activity are strong during working memory tasks, planning, etc. At rest, this activity drops, and internal coherence drops, tending to instead shift to follow patterns in the default mode network ^36^. Our data demonstrates this principle exactly (Figure 4). FC increases significantly during the stress test when cognitive load is high. As the participant relaxes and interacts with a dog, higher-order executive function is no longer needed. Activity within the region and coherence drops accordingly. Of note, FC during the direct interaction phase is also significantly lower than the baseline phase, suggesting a shift toward a lower demand, safety state with reduced need for sustained top down control and self monitoring. This might reflect diminished emotional stress rather than neural disengagement.

**Figure 4:**
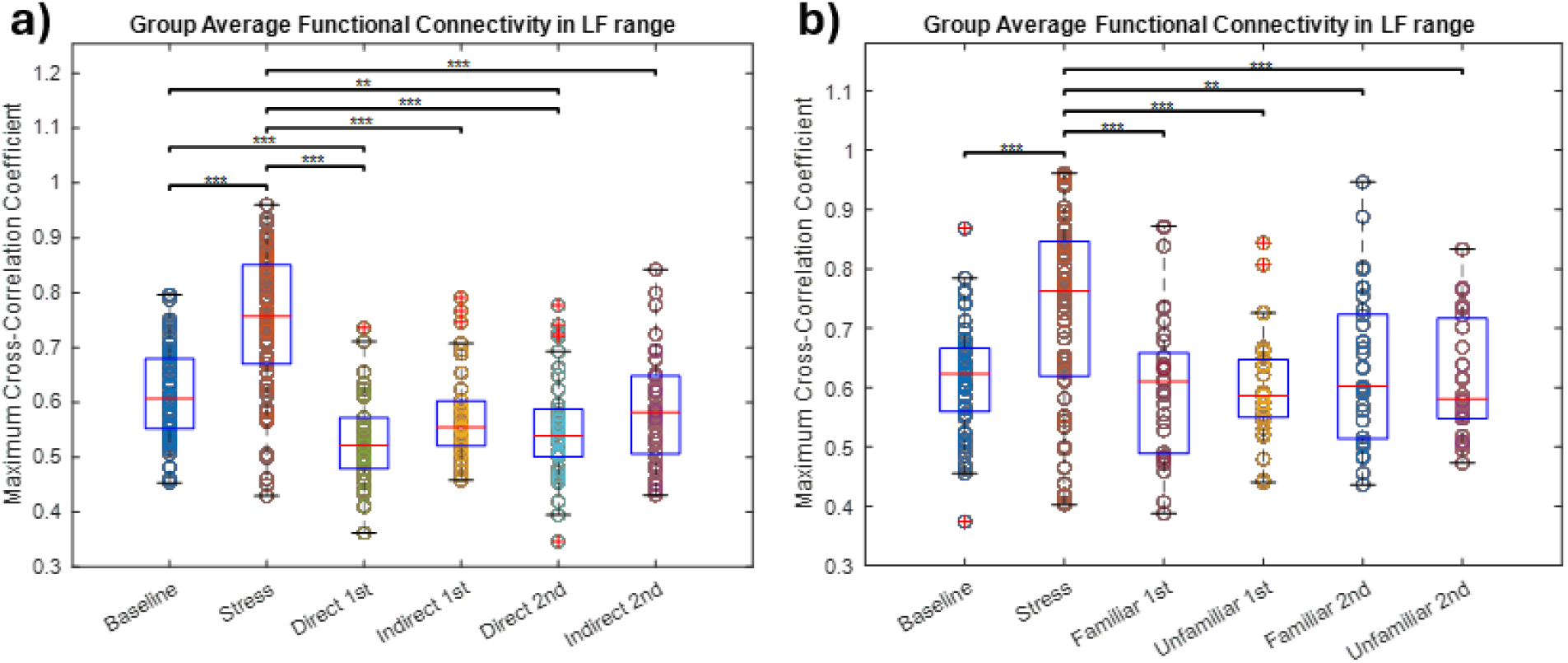
Average Functional Connectivity during Stress Testing (Group A: left column, Group B: right column). A, B) Average maximum cross-correlation coefficient measured at baseline across stress and relaxation phases. Relaxation phase split into 1^st^ and 2^nd^ 10-minute periods. Asterisks indicate significant differences between groups (*: p<0.05, **: p<0.01, ***: p<0.001)

### Perceived Stress and Cortisol

The common stress metrics, VAS and cortisol, were collected at three time points during the study: baseline, post-stress, and post-interaction phase. This study is part of a larger project examining the role of human-animal interaction on acute stress recovery, and specific analyses on VAS and cortisol between interaction type and animal familiarity are presented separately.

To determine if these metrics are related to fNIRS-derived measures, linear regression was performed. Significant interactions were seen for cortisol only with FC in group A. For VAS, significant interactions were observed for all metrics, but only for HRV, LF/HF, and FC across both groups. This might indicate that perceived stress has origins both psychologically, cognitive load in the PFC, and physiologically, the ratio of sympathetic to parasympathetic nervous system activity. This has exciting implications in the field of stress recovery and further demonstrates that dog intervention is a promising tool for stress relief, as HRV, LF/HF, and FC were all significantly affected by all dog conditions.

### Limitations

This study’s participants were largely female. Though recorded, we did not account for dog behavior, i.e., dog excitability, level of interaction with the owner, in these analyses. Future work will examine the interplay between human and dog behavior and physiology.

## Conclusion

This study examined the role of dog intervention in stress recovery from a psychological and physiological perspective. The mode of dog interaction revealed that while all dog interaction types (direct and indirect, unfamiliar dog and familiar dog) were effective in lowering common stress metrics, direct interaction provided the strongest outcomes, outpacing even the resting state. The results from this study can greatly inform the development of future animal-assisted interventions.

## Acknowledgements

The authors would like to thank Rita de Kassia Rodrigues Bezerra Filgueira, the veterinary behavior medicine team, and undergraduate students for their generous assistance in data collection.

## Funding

This study was supported by the Human Animal Bond Research Institute (HABRI) in partnership with Pet Partners, leaders in advancing the science and practice of human-animal interaction (Grant ID: HAB22-025).

## Notes

### Competing Interest Statement

The authors have declared no competing interest.

## References

1. Friedman E, Krause-Parello CA. Companion animals and human health: benefits, challenges, and the road ahead for human–animal interaction. Revue Scientifique et Technique de l’OIE 2018; 37: 71–82.

2. Baun MM, Bergstrom N, Langston NF, et al. Physiological effects of human/companion animal bonding. Nurs Res 1984; 33: 126–9.

3. Barcelos AM, Kargas N, Maltby J, et al. A framework for understanding how activities associated with dog ownership relate to human well-being. Sci Rep 2020; 10: 11363.

4. Lass-Hennemann J, Schäfer SK, Römer S, et al. Therapy Dogs as a Crisis Intervention After Traumatic Events? – An Experimental Study. Front Psychol; 9. Epub ahead of print 4 September 2018. DOI: 10.3389/fpsyg.2018.01627.

5. Bowen J, Bulbena A, Fatjó J. The Value of Companion Dogs as a Source of Social Support for Their Owners: Findings From a Pre-pandemic Representative Sample and a Convenience Sample Obtained During the COVID-19 Lockdown in Spain. Front Psychiatry; 12. Epub ahead of print 14 April 2021. DOI: 10.3389/fpsyt.2021.622060.

6. Weigand KA, Yatcilla JK. Systematic Reviews on Human–Animal Interaction Topics: A Look at Reporting Practices. Anthrozoos 2024; 37: 637–649.

7. Beetz A, Uvnäs-Moberg K, Julius H, et al. Psychosocial and Psychophysiological Effects of Human-Animal Interactions: The Possible Role of Oxytocin. Front Psychol; 3. Epub ahead of print 2012. DOI: 10.3389/fpsyg.2012.00234.

8. Surma S, Oparil S, Narkiewicz K. Pet Ownership and the Risk of Arterial Hypertension and Cardiovascular Disease. Curr Hypertens Rep 2022; 24: 295–302.

9. Rodriguez KE, Delzio MC, Ruekgauer AH, et al. The Effect of a Dog on Acute Stress Reactivity During Experimental Stressors: A Systematic Review and Meta-Analysis. Anthrozoos 2026; 39: 183–209.

10. Polheber JP, Matchock RL. The presence of a dog attenuates cortisol and heart rate in the Trier Social Stress Test compared to human friends. J Behav Med 2014; 37: 860–867.

11. Odendaal JSJ, Meintjes RA. Neurophysiological Correlates of Affiliative Behaviour between Humans and Dogs. The Veterinary Journal 2003; 165: 296–301.

12. Matijczak A, Yates MS, Ruiz MC, et al. The influence of interactions with pet dogs on psychological distress. Emotion 2024; 24: 384–396.

13. Janssens M, Janssens E, Eshuis J, et al. Companion Animals as Buffer against the Impact of Stress on Affect: An Experience Sampling Study. Animals 2021; 11: 2171.

14. Harvie HMK, Rodrigo A, Giuliano RJ. Assessing Whether Household Pets Buffer Responses to a Remote Stress Induction. Anthrozoos 2025; 38: 545–564.

15. Grossberg JM, Alf EF, Vormbrock JK. Does Pet Dog Presence Reduce Human Cardiovascular Responses to Stress? Anthrozoos 1988; 2: 38–44.

16. Gandenberger J, Ledreux A, Taeckens A, et al. The Presence of a Pet Dog Is Associated with a More Balanced Response to a Social Stressor. Stresses 2024; 4: 598–613.

17. Ein N, Reed MJ, Vickers K. Effect of Tranquil and Active Video Representations of an Unfamiliar Dog on Subjective Mental States. Society & Animals 2020; 30: 445–460.

18. Binfet J-T, Green FLL, Draper ZA. The Importance of Client–Canine Contact in Canine-Assisted Interventions: A Randomized Controlled Trial. Anthrozoos 2022; 35: 1–22.

19. Powell L, Edwards KM, Michael S, et al. Effects of Human–Dog Interactions on Salivary Oxytocin Concentrations and Heart Rate Variability: A Four-Condition Cross-Over Trial. Anthrozoos 2020; 33: 37–52.

20. Thayer ER, Stevens JR. Effects of Human-Animal Interactions on Affect and Cognition. Hum Anim Interact Bull. Epub ahead of print 10 October 2022. DOI: 10.1079/hai.2022.0015.

21. Allen K, Blascovich J, Medes W. Cardiovascular Reactivity and the Presence of Pets, Friends, and Spouses: The Truth About Cats and Dogs. Psychosom Med 2002; 64: 727–739.

22. Martos-Montes R, Ordóñez-Pérez D, Ruiz-Maatallah J, et al. Psychophysiological effects of human-dog interaction in university students exposed to a stress-induced situation using the Trier Social Stress Test (TSST). Hum Anim Interact Bull. Epub ahead of print 1 December 2020. DOI: 10.1079/hai.2020.0010.

23. Machová K, Procházková R, Vadroňová M, et al. Effect of Dog Presence on Stress Levels in Students under Psychological Strain: A Pilot Study. Int J Environ Res Public Health 2020; 17: 2286.

24. Marti R, Petignat M, Marcar VL, et al. Effects of contact with a dog on prefrontal brain activation in patients in a minimally conscious state: A controlled crossover trial. Neuroscience 2025; 577: 175–189.

25. Arnskötter W, Marcar VL, Wolf M, et al. Animal presence modulates frontal brain activity of patients in a minimally conscious state: A pilot study. Neuropsychol Rehabil 2022; 32: 1324–1336.

26. Yoo O, Wu Y, Han JS, et al. Psychophysiological and emotional effects of human–Dog interactions by activity type: An electroencephalogram study. PLoS One 2024; 19: e0298384.

27. Lee H-S, Lee J-Y. Assessing the restorative effects of observing a video of dog play in urban dog parks using EEG. Human-Animal Interactions. Epub ahead of print 4 October 2024. DOI: 10.1079/hai.2024.0037.

28. Garrido LFC, Daros RR, Vandresen B, et al. Watch dogs: A mixed-methods investigation of dog owners’views on dog monitoring technologies. Int J Hum Comput Stud 2025; 205: 103645.

29. Smeets T, Cornelisse S, Quaedflieg CWEM, et al. Introducing the Maastricht Acute Stress Test (MAST): A quick and non-invasive approach to elicit robust autonomic and glucocorticoid stress responses. Psychoneuroendocrinology 2012; 37: 1998–2008.

30. Chiarelli AM, Maclin EL, Fabiani M, et al. A kurtosis-based wavelet algorithm for motion artifact correction of fNIRS data. Neuroimage 2015; 112: 128–137.

31. Frasier RM, Starski PA, de Oliveira Sergio T, et al. Sex differences in heart rate variability measures that predict alcohol drinking in rats. Addiction Biology; 29. Epub ahead of print 19 March 2024. DOI: 10.1111/adb.13387.

32. Tong Y, Yao J (Fiona), Chen JJ, et al. The resting-state fMRI arterial signal predicts differential blood transit time through the brain. Journal of Cerebral Blood Flow & Metabolism 2019; 39: 1148–1160.

33. Schaal NK, Hepp P, Schweda A, et al. A Functional Near-Infrared Spectroscopy Study on the Cortical Haemodynamic Responses During the Maastricht Acute Stress Test. Sci Rep 2019; 9: 13459.

34. Schaal K, Schweda A. A functional near-infrared Spectroscopy Study on the cortical Haemodynamic Responses During the Maastricht Acute Stress test. DOI: 10.1038/s41598-019-49826-2.

35. Kim H-G, Cheon E-J, Bai D-S, et al. Stress and Heart Rate Variability: A Meta-Analysis and Review of the Literature. Psychiatry Investig 2018; 15: 235–245.

36. Medvedev A V., Kainerstorfer JM, Borisov S V., et al. Functional connectivity in the prefrontal cortex measured by near-infrared spectroscopy during ultrarapid object recognition. J Biomed Opt 2011; 16: 016008.

